# Differences in social brain function in autism spectrum disorder are linked to the serotonin transporter

**DOI:** 10.1101/2021.05.28.446151

**Authors:** Nichol M.L. Wong, Ottavia Dipasquale, Federico Turkheimer, James L. Findon, Robert H. Wichers, Mihail Dimitrov, Clodagh M. Murphy, Vladimira Stoencheva, Dene M. Robertson, Declan G. Murphy, Eileen Daly, Grainne M. McAlonan

## Abstract

Alterations in the serotonergic control of brain pathways responsible for facial-emotion processing in people with autism spectrum disorder (ASD) may be a target for intervention. However, the molecular underpinnings of autistic-neurotypical serotonergic differences are challenging to access in vivo. Receptor-Enriched Analysis of functional Connectivity by Targets (REACT) has helped define molecular-enriched fMRI brain networks based on a priori information about the spatial distribution of neurochemical systems from available PET templates. Here, we used REACT to estimate the dominant fMRI signal related to the serotonin transporter (5-HTT) distribution during processing of aversive facial expressions of emotion processing in adults with and without ASD. We first predicted a group difference in baseline (placebo) functioning of this system. We next used a single 20 mg oral dose of citalopram, i.e. a serotonin reuptake inhibitor, to test the hypothesis that network activity in people with and without ASD would respond differently to inhibition of 5-HTT. To confirm the specificity of our findings, we also repeated the analysis with 5-HT_1A,_ 5-HT_1B_, 5-HT_2A_, and 5-HT_4_ receptor maps.

We found a baseline group difference in the 5-HTT-enriched response to faces in the ventromedial prefrontal cortex. A single oral dose of citalopram ‘shifted’ the response in the ASD group towards the neurotypical baseline but did not alter response in the control group.

Our findings suggest that the 5HTT-enriched functional network is dynamically different in ASD during processing of socially relevant stimuli. Whether this acute neurobiological response to citalopram in ASD translates to a clinical target will be an important next step.

## Introduction

Autism spectrum disorder (ASD) is neurodevelopmental condition that is characterised by social communication difficulties, in addition to restricted and repetitive behaviours ^1,2^. Altered processing of socio-emotional stimuli may partly contribute to the phenotype. However, there are inconsistencies across the large number of fMRI studies of facial-emotion processing in ASD ^3–10^ and a recent study in 205 individuals with ASD found no significant differences from neurotypical controls ^11^. A possible explanation is that there is simply no common alteration in the neurobiology of social-emotional behaviour which is shared by all individuals on this heterogeneous spectrum ^12^. Alternatively, it is also possible we have not adequately interrogated large-scale brain networks related to the neurochemical systems involved in complex higher-order behaviours ^13^. This may be especially important for the serotonin (5-HT) system which has widespread regulatory effects across brain ^14^, is involved in processing facial emotions ^15–17^, and is strongly implicated in ASD ^18^.

To support this contention, we previously reported preliminary evidence for central serotonin-based anomalies in ASD ^19^, including the functional response to facial expressions of emotion ^8^. Specifically, using fMRI we discovered that lowering 5-HT levels in individuals with ASD using a tryptophan depletion protocol shifts the brain functional response to emotional faces towards a more neurotypical pattern ^8^. We have also confirmed that the manipulation of extracellular 5-HT levels by an acute dose of citalopram, a selective serotonin reuptake inhibitor (SSRI), alters the dynamics of the functional response to emotional faces in individuals with ASD, but not in neurotypicals, in the core social brain regions including amygdala, ventromedial prefrontal cortex (vmPFC), and striatum ^20^. Taken together this suggests that the brain’s 5-HT system is dynamically different in ASD in response to emotional faces, but we have yet to establish its molecular underpinnings.

The 5-HT transporter (5-HTT) is an integral plasma membrane protein that actively transports 5-HT for clearance and regulation of extracellular 5-HT. It is critical for 5-HT neurotransmission and homeostasis. Also, animal studies reported that 5-HTT knockout mice have autism-like behavioural phenotypes ^21^. Moreover, a genetic study of the 5-HTT locus SLC6A4 identified multiple rare variants of 5-HTT with significant linkage to ASD ^22^, albeit this finding could not be replicated in another group ^23^. Finally, some ^24–26^ (but not all ^27^) human molecular imaging studies reported lower availability of regional 5-HTT in individuals with ASD that are related to their difficulties in social functioning. These prior studies were important first steps; however, a potentially crucial question has not been addressed – if the 5-HTT system in ASD *functions* differently.

Therefore, to understand if functional differences associated with the 5-HTT functional network contribute to altered processing of facial emotion in ASD, we employed the Receptor-Enriched Analysis of functional Connectivity by Targets (REACT) approach ^28^. This method allowed us to weight fMRI analysis by the molecular distribution of 5-HTT in the brain, in order to better characterise the fMRI response to pharmacological manipulation of 5-HT during processing of facial expressions of emotion ^28^. Hence, to test the hypothesis that there is a difference in adults with and without ASD, we used a 5-HTT PET template ^29^ as a spatial regressor in a dual-regression analysis ^30^.

In brief, on one of two study visits, either 20 mg citalopram or placebo was administered orally to individuals with and without ASD in a double-blind, randomised order, 3 hours prior to an aversive facial-emotion processing fMRI task. In line with the literature and the mean 5-HTT-enriched functional map resulting from the analysis with REACT during the placebo condition, our functional network of interest comprised key social brain regions (i.e., amygdala, vmPFC, striatum and fusiform gyrus) ^31^ involved in facial emotion processing.

However, 5-HT reuptake inhibition could also have secondary effects through constituent receptors of the serotonergic system (Riad *et al*., 2001; Selvaraj *et al*., 2014; Jackson and Moghaddam, 2018). Hence, to confirm the specificity of our findings, we also conducted supplementary analyses exploring also the 5-HT_1A_-, 5-HT_1B_-, 5-HT_2A_-, and 5-HT_4_- enriched functional networks.

## Materials and Methods

### Participants

Forty right-handed adult males (19 with ASD and 21 neurotypical controls, 18-60 years old) were enrolled in this study. Individuals with a clinical diagnosis of ASD were recruited through the National Adult Autism Service at the Maudsley Hospital where diagnosis is a multi-disciplinary decision supported by the Autism Diagnostic Interview – Revised (where an informant is available) and/or the Autism Diagnostic Observation Schedule (ADOS) ^35^. Only individuals of IQ > 70 with no history of medical disorders that influence cognitive performance, no other major mental illnesses, genetic disorders associated with ASD, no alcohol or substance dependence, and not on any medication affecting the 5-HT system were included. Their scores on anxiety and depression were also measured by Hamilton Anxiety Rating Scale (HAM-A) and Hamilton Depression Rating Scale (HAM-D) respectively ^36,37^ since these symptoms are common in ASD. All participants gave written, informed consent and this study received National Research Ethics approval from the Stanmore NHS Research Ethics Committee (reference: 14/LO/0663).

### Study design

This study adopted a placebo-controlled, randomised, double-blind, repeated-measures, cross-over, case-control study design as part of a larger investigation into the brain response to 5-HT medications in ASD (https://clinicaltrials.gov Identifier: NCT04145076). Placebo and citalopram were allocated in a pseudo-randomised order, approximately half in each group attended a placebo visit before citalopram and the other half attended a citalopram visit before placebo, generated using an algorithm (https://www.random.org/). Participants completed two scanning visits that were separated by at least eight days, and they were administered an acute dose of encapsulated citalopram (20mg) or of encapsulated placebo (ascorbic acid) three hours before each scanning session as based on the pharmacokinetic properties of citalopram ^38^.

During each scanning session, a face-emotion processing task ^20,39^ was administered to participants. Briefly, the task consisted of four blocks of faces and four blocks of geometric shapes, with six five-second trials per block. Participants had to indicate which of the two images in the lower panel was identical to the target image in the upper panel in each trial. The facial images were either ‘angry’ or ‘fearful’. Full details of the study design have been reported elsewhere ^20^.

### MRI data acquisition and preprocessing

The fMRI data were acquired on a 3T General Electric Signa HD × Twinspeed scanner (Milwaukee, Wisc.) fitted with a quadrature birdcage head coil: TR = 2,000 ms, TE = 30 ms, FOV = 192 × 192 mm, voxel size = 3 × 3 × 5 mm, flip angle = 80°, number of time points = 135. T1-weighted structural data was acquired in the sagittal planes: TR = 7.312 ms, TE = 3.016 ms, FOV = 270 × 270 mm^2^, matrix size = 256 × 256, slice thickness = 1.2 mm, flip angle = 11°.

Each participant’s fMRI images were first corrected for slice-timing and head movement with *FSL* ^40^. Participants with head movement of ≥ 3 mm in any plane, rotation ≥ 3° in any direction, or with > 30% of their timepoints having framewise displacement (FD) > 0.2 mm ^41^ were excluded, giving us a final sample of 21 controls and 13 individuals with ASD for subsequent analyses. A temporal high-pass filter at 128 s was applied and the fMRI data were smoothed by 8 mm full width at half-maximum kernel, normalised to the MNI space using their skull-stripped structural brain images and resampled to 2 mm^3^.

### 5-HT-enriched functional response during task

For the following analysis, we used the high-resolution in-vivo atlas of the distribution density of 5-HTT ^29^, that is characterised by high intensity in the entorhinal and insular cortices, subcortical regions, and the brainstem (**Figure 1 A)**.

**Figure 1.**
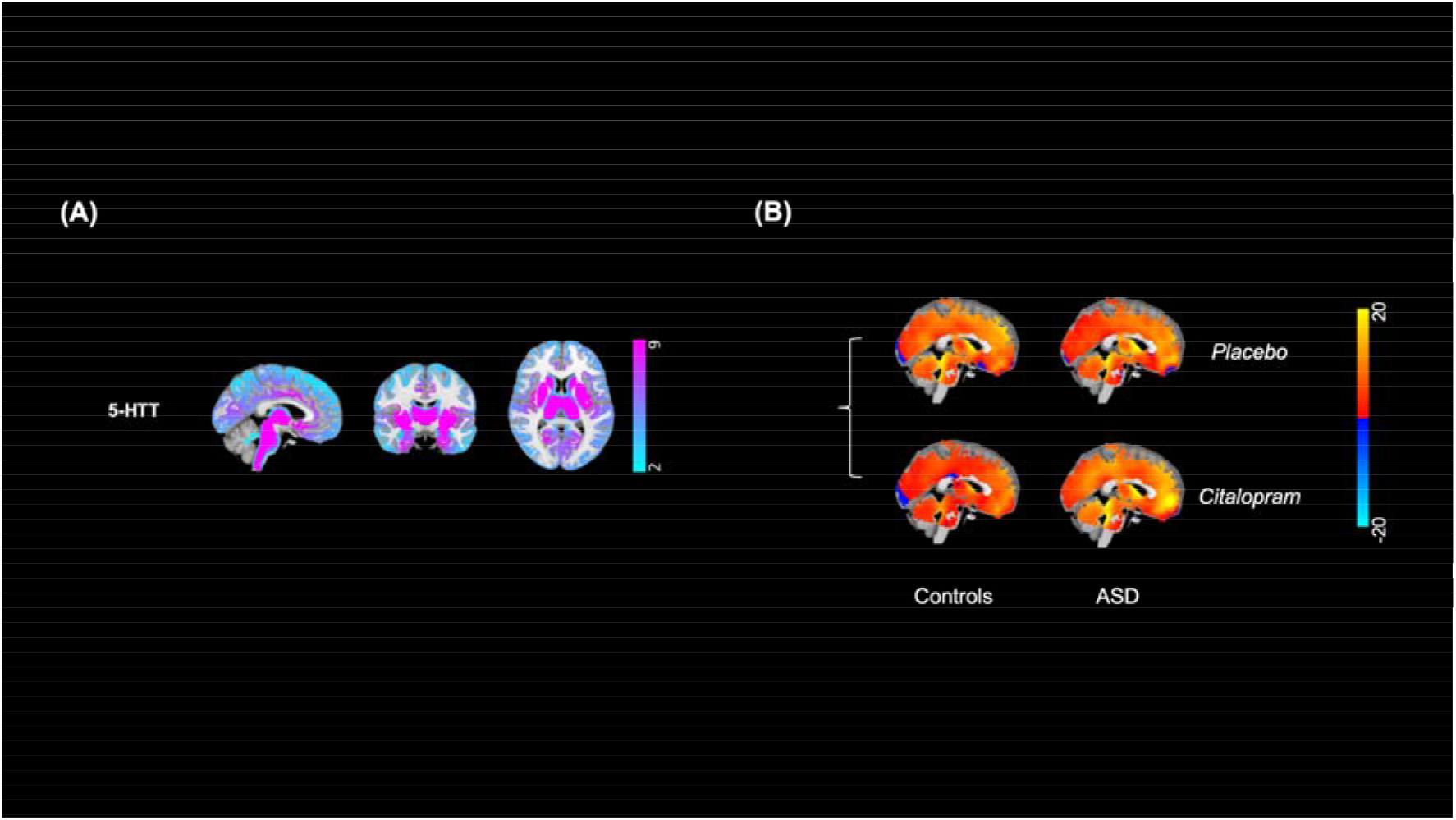
(**A**) The in-vivo atlas of the serotonin transporter (5-HT_T_) is presented with its corresponding density distribution on a colour scale. (**B**) The 5-HTT-enriched functional maps corresponding to the interaction between the convolved Faces>Shapes regressor and the dominant blood-oxygen-level-dependent (BOLD) fluctuation within 5-HTT map were averaged in individuals with and without autism spectrum disorder (ASD) in each drug condition for visualisation.

The atlas was used as a molecular template to estimate the 5-HTT-enriched functional response during the task in a two-step regression using the REACT approach ^28^ with *FSL* ^40^. Details of this methodology can be found elsewhere ^28^. In brief, in the first step, the molecular template is used as a spatial regressor to weight the fMRI signal in the grey matter (for this step the images were resampled at 1 mm^3^) and estimate the dominant blood-oxygen-level-dependent (BOLD) fluctuation of the 5-HTT-enriched functional system at the subject level. The cerebellum was excluded at this stage as it was used as a reference region in the kinetic model for the estimation of the 5-HTT PET atlas ^29^. In the second step, the resulting subject-specific time-series were incorporated as a temporal regressor of 5-HTT functional system into a generalised psycho-physiological interaction (gPPI) design to estimate the 5-HTT-enriched Faces>Shapes functional response of the brain (including the cerebellum; resampled at 2 mm^3^). Both data and design were demeaned in both steps for each subject; the design matrix was also normalised to the unit standard deviation in the second step. The design in the second step included the convolved Faces>Shapes regressor and its derivative, the convolved regressor with both Faces and Shapes blocks and its derivative, the dominant BOLD fluctuation within the 5-HTT map, the regressor of the interaction between the convolved Faces>Shapes regressor and this BOLD fluctuation, six translational and rotational parameters of head movement, and the temporal masking regressor(s) of timepoints with FD > 0.2 mm. Our regressor of interest was the regressor of interaction between the convolved Faces>Shapes regressor and the dominant BOLD fluctuation within the 5-HTT map. The averaged 5-HTT-enriched BOLD functional response across participants is visualised in **Figure 1 B**.

We also explored the functional response with a separate design that incorporated four more dominant BOLD fluctuation regressors within other 5-HT spatial PET maps (i.e., 5-HT_1A_, 5-HT_1B_, 5-HT_2A_, 5-HT_4_) in addition to the dominant BOLD fluctuation within the 5-HTT map in the same model and the results are included in the **Supplementary Information**.

### Statistical analyses

Demographics and scores in anxiety and depression were compared between groups using t-tests. Task behavioural performances in terms of response accuracy and reaction time (RT) between groups, stimuli, and drug conditions were compared using analysis of covariance (ANCOVA), controlling for their scores in anxiety and depression because of the group differences. Analyses were performed in *R* (https://www.r-project.org) and significance was inferred when p < 0.05.

In the task functional response analyses, whole brain voxel-wise analysis was first performed in placebo condition to identify the baseline Faces>Shapes 5-HTT-enriched functional response across individuals with and without ASD using general linear modelling (and controlling for their scores in anxiety and depression). We next compared the 5-HTT-enriched Faces>Shapes functional response during placebo and citalopram conditions within our regions of interest (ROIs) – amygdala, vmPFC, striatum, and fusiform gyrus – between each group using *randomise* in *FSL* ^42^. The ROIs were derived from Juelich histological atlas, vmPFC atlas of asymmetric and probabilistic cytoarchitectonic maps, Harvard-Oxford subcortical structural atlas and anatomical probability maps ^43–45^. Given the group differences in anxiety and depression ratings, these scores were also entered as covariates. Significance was inferred when family-wise-error (FWE) corrected p_FWE_ < 0.05 using permutation testing with 5000 permutations and threshold-free cluster enhancement ^46^.

Finally, with the caveat that this study was not powered to investigate multiple clinical measures, we carried out a preliminary exploration of whether an autistic individual’s age, IQ, or ADOS subscores were related to their 5-HTT-enriched functional response using multiple linear regression. Significance was inferred when p < 0.05.

## Results

### Demographics and clinical characteristics

Individuals with and without ASD did not differ significantly in terms of age (t_31_ = 0.892, p = 0.379) and IQ (t_31_ = 0.344, p = 0.733). As expected, however, individuals with ASD had higher scores in anxiety (t_31_ = 3.298, p = 0.002) and depression (t_19.2_ = 2.900, p = 0.009) ratings (**Table 1**).

**Table 1.** Demographics and clinical characteristics of the study sample

|  | **ASD**<br>(n=13) | **Controls**<br>(n=21) | ***T*** | ***p*** |
| --- | --- | --- | --- | --- |
| **Demographics** |  |  |  |  |
| *Age (years)* | 30.31 (11.54) | 27.05 (9.34) | 0.892 | 0.379 |
| *Intelligence quotient* | 112.77 (18.36) | 114.45 (9.73) | 0.344 | 0.733 |
| **Clinical characteristics** |  |  |  |  |
| *HAM-A* | 8 (5.05) | 3 (4.18) | 3.298 | 0.002 |
| *HAM-D* | 6 (4.65) | 2 (3.16) | 2.900 | 0.009 |
| *ADOS-C* | 3 (2.02) | - |  |  |
| *ADOS-S* | 6 (2.63) | - |  |  |

The means are presented with standard deviations in parenthesis.
HAM-A=Hamilton Anxiety Rating Scale; HAM-D=Hamilton Depression Rating Scale;
ADOS-C=Autism Diagnostic Observation Schedule–Communication domain;
ADOS-S=Autism Diagnostic Observation Schedule–Reciprocal Social Interaction domain

### Task behavioural performance

The accuracy of the individual responses during the task for the two drug conditions was investigated using a group × drug × stimuli factorial design and no interaction effects were identified (p > 0.05) (**Table 2**). There was only a significant main effect of stimuli type, with individuals having higher accuracy in matching faces than shapes regardless of group (F_1,28_ = 23.27, p < 0.001). Regarding response reaction times, no interaction effects were identified (p > 0.05) but there was a significant main effect of stimuli type - with individuals matching faces more slowly than shapes regardless of group (F_1,28_ = 15.14, p = 0.001) (**Table 2**).

**Table 2.** Behavioural performance of the study sample during the task

|  | Stimuli | Drug | Group | Mean (SD) |
| --- | --- | --- | --- | --- |
| <u>Accuracy (%):</u> |  |  |  |  |
|  | Shapes | Placebo | Controls | 93.20 (6.55) |
|  |  |  | ASD | 94.87 (5.68) |
|  |  | Citalopram | Controls | 94.74 (5.17) |
|  |  |  | ASD | 94.55 (4.93) |
|  | Faces | Placebo | Controls | 97.81 (5.95) |
|  |  |  | ASD | 99.36 (1.57) |
|  |  | Citalopram | Controls | 98.90 (3.06) |
|  |  |  | ASD | 99.36 (1.57) |
| <u>Reaction time (seconds):</u> |  |  |  |  |
|  | Shapes | Placebo | Controls | 0.98 (0.20) |
|  |  |  | ASD | 1.13 (0.30) |
|  |  | Citalopram | Controls | 1.00 (0.24) |
|  |  |  | ASD | 1.10 (0.31) |
|  | Faces | Placebo | Controls | 1.04 (0.21) |
|  |  |  | ASD | 1.23 (0.27) |
|  |  | Citalopram | Controls | 1.09 (0.23) |
|  |  |  | ASD | 1.23 (0.30) |

The means and standard deviations (SD) are presented.

### Baseline placebo 5-HTT-enriched functional response

Using whole brain voxel-wise analysis, a significant increase in 5-HTT-enriched Faces>Shapes response was identified at baseline placebo condition across individuals with and without ASD within regions including the superior temporal gyrus (STG), superior parietal lobule (SPL), posterior cingulate cortex (PCC), amygdala, vmPFC, striatum, and fusiform gyrus, confirming that the 5-HTT-enriched functional response was within our hypothesised ROIs (p_FWE_ < 0.05) (**Figure 2; Table 3**).

**Figure 2.**
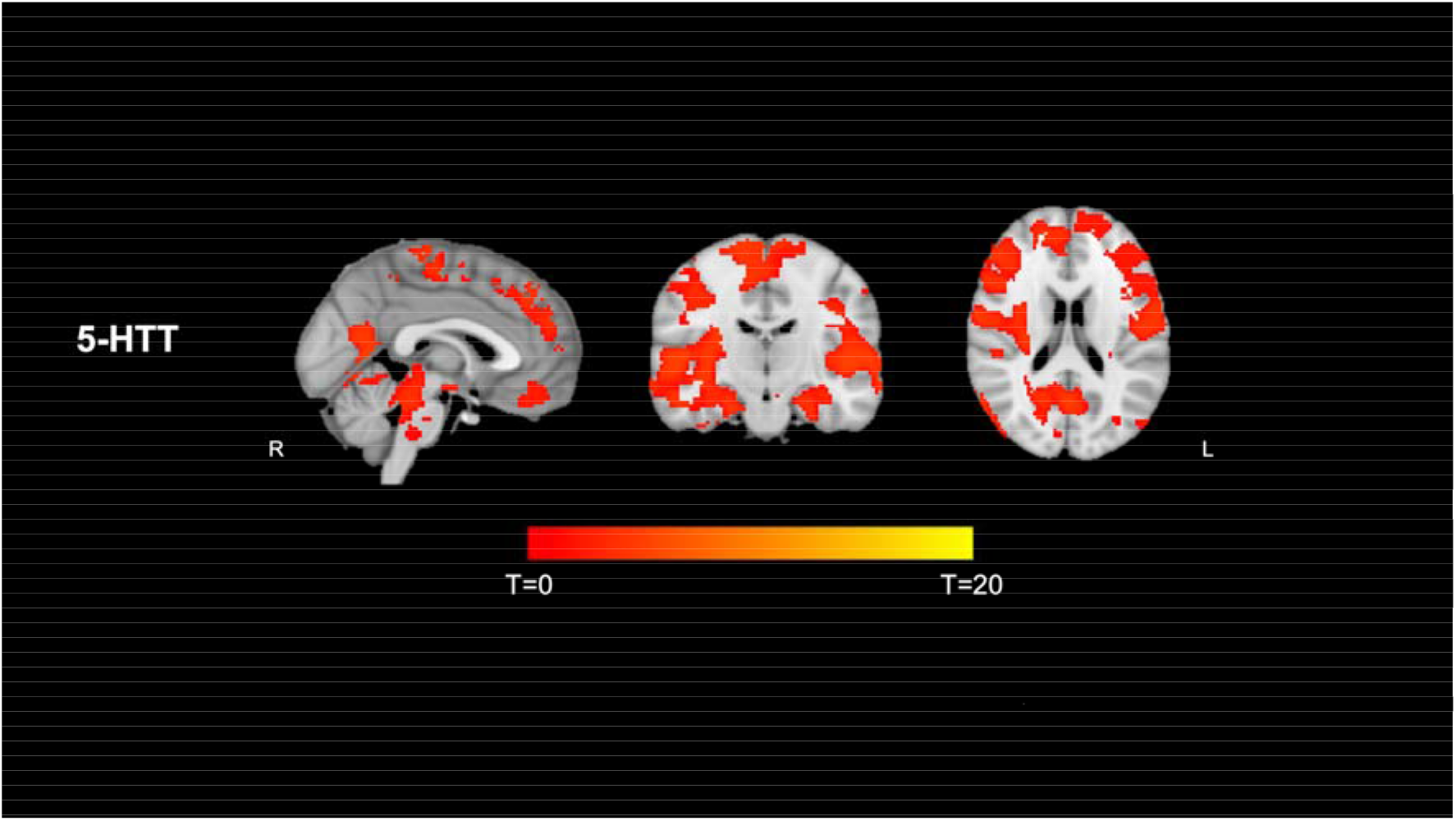
The significant clusters of increased serotonin transporter (5-HTT)-enriched Faces>Shapes functional response across individuals with and without autism spectrum disorder (ASD) are overlaid on the anatomical brain. L=Left, R=Right.

**Table 3.** The clusters with increased 5-HTT-enriched Faces>Shapes functional response at baseline placebo condition across the whole sample

| Cluster | Size (voxels) | MNI coordinates |  |  | **P<sub>peak</sub>** |
| --- | --- | --- | --- | --- | --- |
|  |  | *X* | *Y* | *Z* |  |
| **1** | 48525 | 54 | -16 | 2 | 0.037 |
| **2** | 379 | -28 | -52 | 34 | 0.049 |
| **3** | 70 | 10 | -38 | 26 | 0.050 |
| **4** | 69 | -28 | -32 | 18 | 0.050 |

### Group comparisons of the 5-HTT-enriched functional response

To investigate any differences in the action of citalopram in individuals with and without ASD, we performed an voxel-wise analysis on the 5-HTT-enriched functional connectivity within specific ROIs. At the baseline placebo condition, we observed a significant group difference (Controls>ASD) in the 5-HTT-enriched functional response in the right vmPFC (specifically within the medial orbitofrontal cortex [mOFC] and subgenual anterior cingulate cortex [ACC]) (k = 70, p_peak_ = 0.039, MNI_xyz_ = [16, 22, -18]); in contrast, there was no ASD-Controls group difference in the citalopram condition. Although we did not find any significant group × drug interaction effect across voxels, post-hoc analyses revealed that citalopram increased the 5-HTT-enriched functional response in ASD (p = 0.018) to a response level similar to the neurotypical controls (p > 0.05). There was no significant response difference between baseline placebo and citalopram in neurotypical controls (p = 0.541) (**Figure 3**). To confirm the specificity of the findings, we repeated the analysis by including all *five* dominant BOLD fluctuations of the 5-HT systems in the same model and found similar differences in 5-HTT-enriched functional response, suggesting that the response observed is specific to dominant BOLD fluctuations within 5-HTT map and could not be explained by other 5-HT receptors. Details are reported in the **Supplementary Information**.

**Figure 3.**
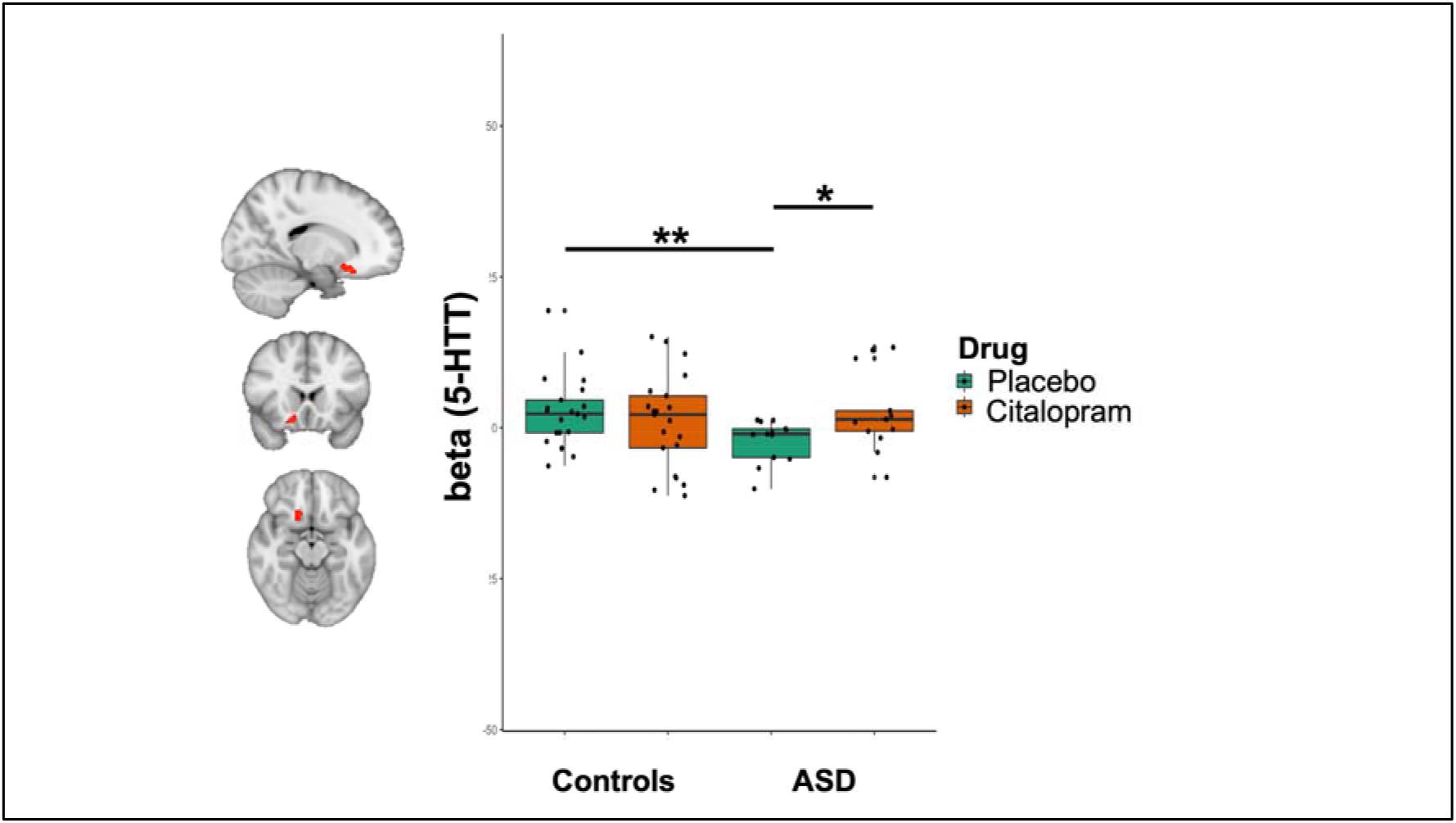
Significantly higher serotonin transporter (5-HTT)-enriched Faces>Shapes response in neurotypical controls compared to individuals with autism spectrum disorder (ASD) within the right ventromedial prefrontal cortex (vmPFC). Post-hoc analysis revealed that citalopram significantly increased the 5-HTT-enriched functional response in ASD. The results are demonstrated in a box plot. *p<0.05, **p<0.01.

### Individual variations and 5-HTT-enriched functional response in ASD

We also explored whether age, IQ, and clinical characteristics might be related to the citalopram-induced changes in 5-HTT-enriched functional response in individuals with ASD. Citalopram caused the greatest change in the mean functional response across the right subgenual ACC (ß = -0.697, p = 0.034) in autistic individuals with the least communication difficulties. Age, IQ, scores of anxiety and depression were not related to citalopram-induced changes in 5-HTT-enriched functional response in ASD, suggesting that the functional differences in the 5-HTT social brain network are not due to co-occurring mental health difficulties or overall intellectual ability.

## Discussion

We found a baseline ASD-neurotypical group difference in the function of a ‘social brain’ 5-HTT network. However, following a single oral dose of citalopram (which inhibits the 5-HTT), the response within vmPFC in the ASD group was no longer different from the neurotypical baseline (placebo). Thus, the functional dynamics of 5-HTT pathways involved in facial emotion processing are altered in ASD but can be ‘shifted’ to a neurotypical placebo profile by inhibition of 5-HTT.

Specifically, face-emotion processing activated a 5-HTT-enriched functional circuit across the STG, PCC, amygdala, vmPFC, striatum, and fusiform gyrus, i.e. regions which underpin face and facial emotion processing ^31^. However, we observed a significantly lower response to emotional faces in the mOFC and subgenual ACC within the vmPFC in individuals with ASD compared to neurotypical controls at baseline (placebo). The medial prefrontal cortex (mPFC) is crucial to the theory of mind, which is at least partly dependent upon facial-emotion processing ^47^. Therefore, our placebo result converges with prior studies that reported altered vmPFC response during processing of facial emotions in ASD ^5,10^, but extends to support a critical role for 5-HTT in ASD.

Our results reveal a potential functional difference in 5-HTT during facial emotion processing. This was confirmed by the effect of citalopram which acts on 5-HTT (in addition to other 5HT receptors) and which increased the vmPFC response in individuals with ASD to a similar level to that observed in neurotypical adults. In contrast, there was no change in brain function in neurotypical adults as measured 3 hours following the 5-HT challenge by the acute citalopram dose. This shift of functional response in individuals with ASD to a more neurotypical pattern was broadly similar to that observed in our previous study of tryptophan depletion in people with and without ASD ^8^. A tryptophan depletion protocol lowers 5-HT levels and shifts the ASD individuals’ response to fearful and sad faces within regions including mPFC to a neurotypical pattern ^8^. At first glance it is challenging to reconcile the similar actions of 5-HT reduction through acute tryptophan depletion and 5-HTT inhibition by citalopram, which should increase 5-HT levels, in ASD. However, recent re-appraisal of the action of acute tryptophan depletion has concluded that there is “no convincing evidence” that the procedure alters central 5-HT release and/or neuronal activity ^48^. Indeed, the effect of altering tryptophan levels on 5-HT levels “depends critically on the circumstances in which it is given” ^49^. An acute citalopram dose has previously been shown to be similar to the effect of acute tryptophan depletion in behavioural tasks ^50,51^. Hence, to probe the 5-HT system, citalopram is much more specific as it has high affinity for 5-HTT ^52^. This adds confidence that our findings are best explained by a functional difference in the 5-HTT system in ASD.

How functional differences in the 5-HTT arise in ASD is not clear. The 5-HT system is essential for brain development and is also vulnerable to multiple influences during the critical antenatal period. For example, foetal exposure to pharmacologic (e.g. 5-HT reuptake inhibitors) and/or genetic (functional polymorphisms) differences in 5-HTT function (in mother or foetus) are linked with ASD and related neurodevelopmental outcomes ^53,54^. Consequently, inhibiting the 5-HTT activity with SSRIs has been reported to be a useful intervention in children with such complex neurodevelopmental conditions ^55^. The most recent PET investigation of 5-HTT in ASD ^26^ reported lower availability across the brain, but especially in the ACC; lower 5-HTT availability in ACC was also correlated to the extent of difficulties in social functioning ^25^. It is possible that lower levels of a target for citalopram may mean the response to 20 mg dose is easier to detect; i.e. there is less redundancy in the system. This also chimes with evidence that people with ASD are more sensitive to the side effects of SSRI ^56^ and advice to start SSRIs at a lower dose in ASD^55^.

It is noteworthy that the response to citalopram observed in ASD here could not be attributed to the presence of common co-occurring conditions (i.e., anxiety and depression) or current use of medicines impacting on 5-HT. Instead, we found that symptoms on the autism spectrum - mainly on the communication domain - were related to the shift in 5-HTT-enriched functional response in the vmPFC. We observed that individuals with ASD who had less communication difficulties had a greater shift in response to citalopram challenge. In addition, scrutiny of the individual responses indicates that not everyone in the ASD group responded to citalopram in the same direction. This may have important clinical applications as it indicates that not every individual on the heterogeneous autism spectrum will respond in the same way to 5-HT treatment. It lends support to the current guidelines that low dose of SSRI should be used in individuals with ASD, titrating up gradually with careful monitoring ^57^.

## Limitations

There are several limitations of this study. The analyses were based on a relatively small sample size because we adhered to strict recruitment criteria and applied strict motion correction criteria. However, a strength of our study was the repeated visits design which increased power by reducing heterogeneity of the sample. Our hypothesis was limited to the 5-HTT. Future studies could also test the functional response in relation to other receptor density maps, such as GABA and glutamate, which also influence brain function and are modulated by 5-HT ^14^. We did not examine a more general impact of 5-HT modulation on cognitive and affective functioning ^58^. Therefore, caution is warranted before generalising the findings to other processes such as cognitive inhibition. This multimodal REACT approach offers specific insight into the functional response enriched by the 5-HTT system. However, it is noteworthy that the PET molecular templates used here were from a study with typically developed controls ^29^ and although recent evidence suggests lower availability of 5-HTT in ASD ^24–26^, there is disagreement ^27^ and we cannot say what the exact pattern of 5-HTT distribution was in either controls or ASD participants in this study. Other limitations should also be noted, including the use of single-sex sample in order to increase sample homogeneity and to avoid inadvertent exposure of pregnant females to SSRI ^20^. Hence, it is important to extend our work in future to include females with ASD.

## Conclusions

This is the first study to apply the multimodal REACT approach to a task-fMRI paradigm to enrich the analysis by accounting for the distribution of 5-HTT, main target of citalopram. This provided a novel insight into the functional dynamics of the 5-HTT system in ASD during social information processing. An ASD-neurotypical group difference in functional response to facial emotions in the 5-HTT-related functional circuit was found, but disappeared following citalopram administration that leads to 5-HTT inhibition. The response to citalopram was observed only in the ASD group and suggests that the 5-HTT is dynamically different during processing of salient social stimuli in ASD. This may constitute a treatment target which should be explored in future clinical studies.

## Acknowledgement

The authors would like to thank all the volunteers for their participation.

The authors acknowledge support from the Maudsley Pharmacy Department and the National Autism Service for Adults at the South London, the National Institute for Health Research (NIHR) Biomedical Research Centre for Mental Health at South London and Maudsley NHS Foundation Trust and Institute of Psychiatry, Psychology and Neuroscience King’s College London. The views expressed are those of the author(s) and not necessarily those of the IMI, NHS, the NIHR or the Department of Health. Support has also been received funding from the Sackler Institute for Translational Neurodevelopment at King’s College London, the MRC Centre for Neurodevelopmental Disorders at King’s College London and the Innovative Medicines Initiative 2 Joint Undertaking under grant agreement No 777394 for the project AIMS-2-TRIALS. This Joint Undertaking receives support from the European Union’s Horizon 2020 research and innovation programme and EFPIA and AUTISM SPEAKS, Autistica, SFARI.

