## Supplementary Information for "Differences in social brain function in autism spectrum disorder are linked to the serotonin transporter"

***Group comparison of head motion***

Participants with head movement of ≥ 3 mm in any plane, rotation ≥ 3° in any direction, or with > 30% of their timepoints having framewise displacement (FD) > 0.2 mm were excluded for analyses and we have compared the FD in our final sample of 21 controls and 13 individuals with analysis of variance (ANOVA). Our results revealed that there were no significant group × drug (F_1,32_ = 0.601, p = 0.444), and group (F_1,32_ = 1.099, p = 0.302) effects, but significant drug (F_1,32_ = 4.483, p = 0.042) effect on FD in our final sample, suggesting that participants had more head motion after citalopram than baseline placebo (**Figure S1**).

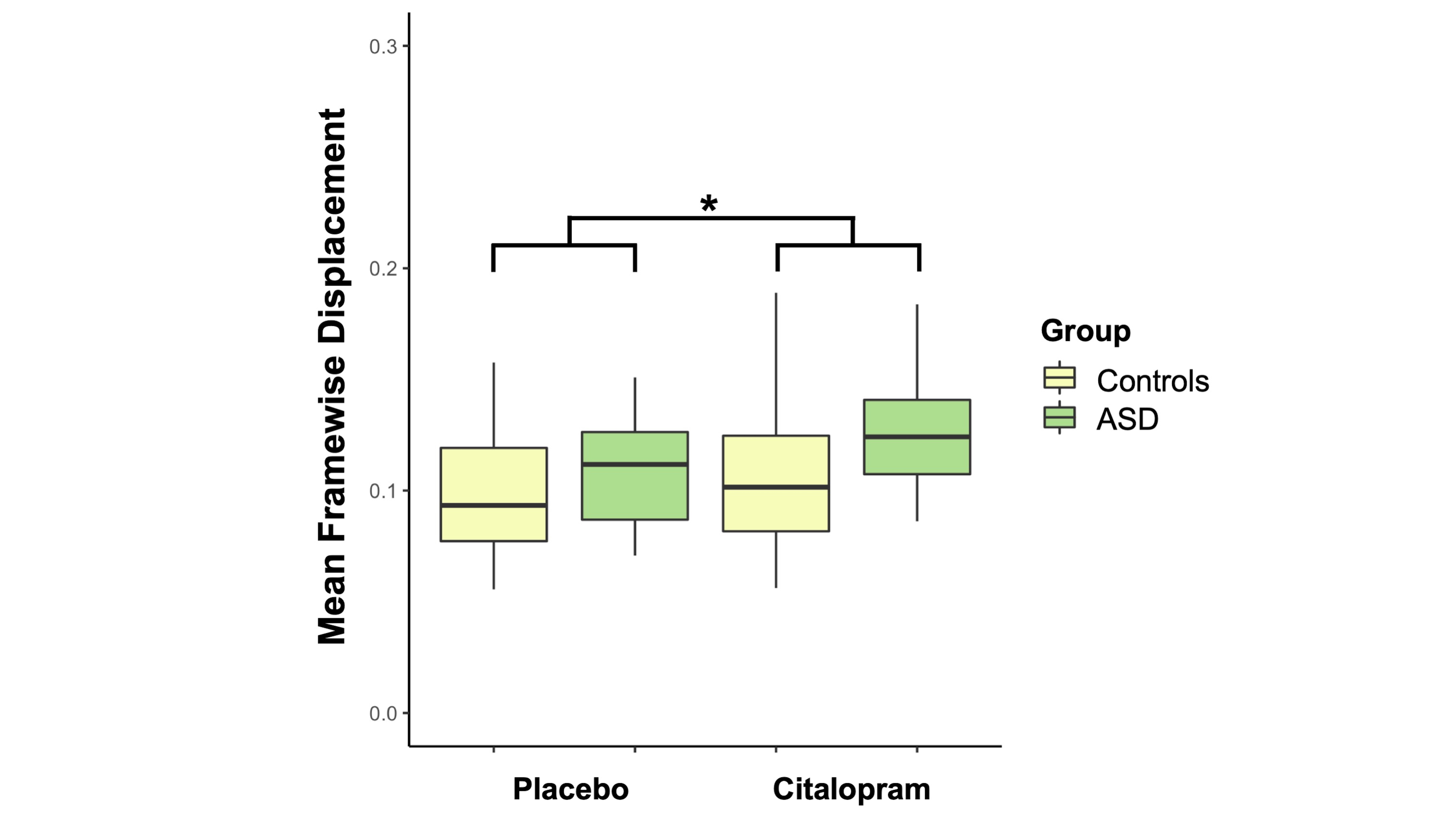

**Figure S1**. The mean framewise displacement (FD) in baseline placebo and citalopram conditions of individuals with and without autism spectrum disorder (ASD) is visualised in a box plot. Analysis of variance (ANOVA) suggested that there was a significant drug effect (p = 0.042) of higher mean FD in citalopram than baseline placebo condition across the whole sample.

***Task behavioural performance and 5-HTT-enriched functional response in ASD***

Although there were no significant drug and group × drug effects on the accuracy and reaction time of the individuals’ responses during the task, we have also explored whether the citalopram-induced change in responses’ accuracy and reaction time might be associated to the change in 5-HTT-enriched functional response in the mOFC and subgenual ACC within the right vmPFC in individuals with ASD based on our group comparisons’ findings. Although there were no group differences in behavioural performance between individuals with and without ASD, we found that the citalopram-induced change in accuracy difference (Faces>Shapes) in individuals with ASD were negatively related to their change in 5-HTT-enriched functional response in the right anterior mOFC (ρ = -0.625, p = 0.022), right posterior mOFC (ρ ≤ -0.704, p ≤ 0.007), and right subgenual ACC (ρ = -0.603, p = 0.029). We did not observe significant associations between the change in reaction time difference (Faces>Shapes) and the change in 5-HTT-enriched response in individuals with ASD.

***Group comparisons of the five 5-HT-enriched functional circuits***

We have also repeated the comparison between individuals with and without ASD their all *five* 5-HT-enriched Faces>Shapes functional response (using 5-HT_1A_, 5-HT_1B_, 5-HT_2A_, 5-HT_4_, and 5-HTT PET maps) within and between placebo and citalopram conditions, controlling for the scores in anxiety and depression, to investigate whether there was an ASD-neurotypical group difference in the functional response within our hypothesised ROIs (amygdala, vmPFC, striatum, and fusiform gyrus). With the same REACT approach, we have included all five dominant BOLD fluctuation regressors within the 5-HT spatial maps. Hence, this additional design in the second step of the dual-regression included the convolved Faces>Shapes regressor and its derivative, the convolved regressor with both Faces and Shapes blocks and its derivative, the *five* dominant BOLD fluctuation regressors within 5-HT_1A_, 5-HT_1B_, 5-HT_2A_, 5-HT_4_ and 5-HTT maps, the regressors of the interaction between the convolved Faces>Shapes regressor and the *five* regressors of 5-HT, six translational and rotational parameters of head movement, and the temporal masking regressors of timepoints with FD > 0.2 mm. Here our regressors of interest were the *five* regressors of interaction between the convolved Faces>Shapes regressor and the regressors of 5-HT.

We found significant group difference (Controls > ASD) in both the 5-HT_1B_-enriched functional response and the 5-HTT-enriched functional response in the baseline placebo condition, but no group difference in the citalopram condition (**Table S1; Figure S2**). We did not find any significant group × drug interaction effect across voxels.

| **Table S1.** The clusters of significant group differences (Controls>ASD) in 5-HT_1B_-enriched and 5-HTT-enriched Faces>Shapes functional response in baseline placebo condition | | | | | |
| --- | --- | --- | --- | --- | --- |
| **Cluster** | **Size (voxels)** | **MNI coordinates** | | | **p_peak_** |
|  |  | ***X*** | ***Y*** | ***Z*** |  |
| **5-HT_1B_:** |  |  |  |  |  |
| 1 | 1974 | 14 | 52 | -2 | 0.014 |
| **5-HTT:** |  |  |  |  |  |
| 1 | 18 | 12 | 40 | -24 | 0.041 |

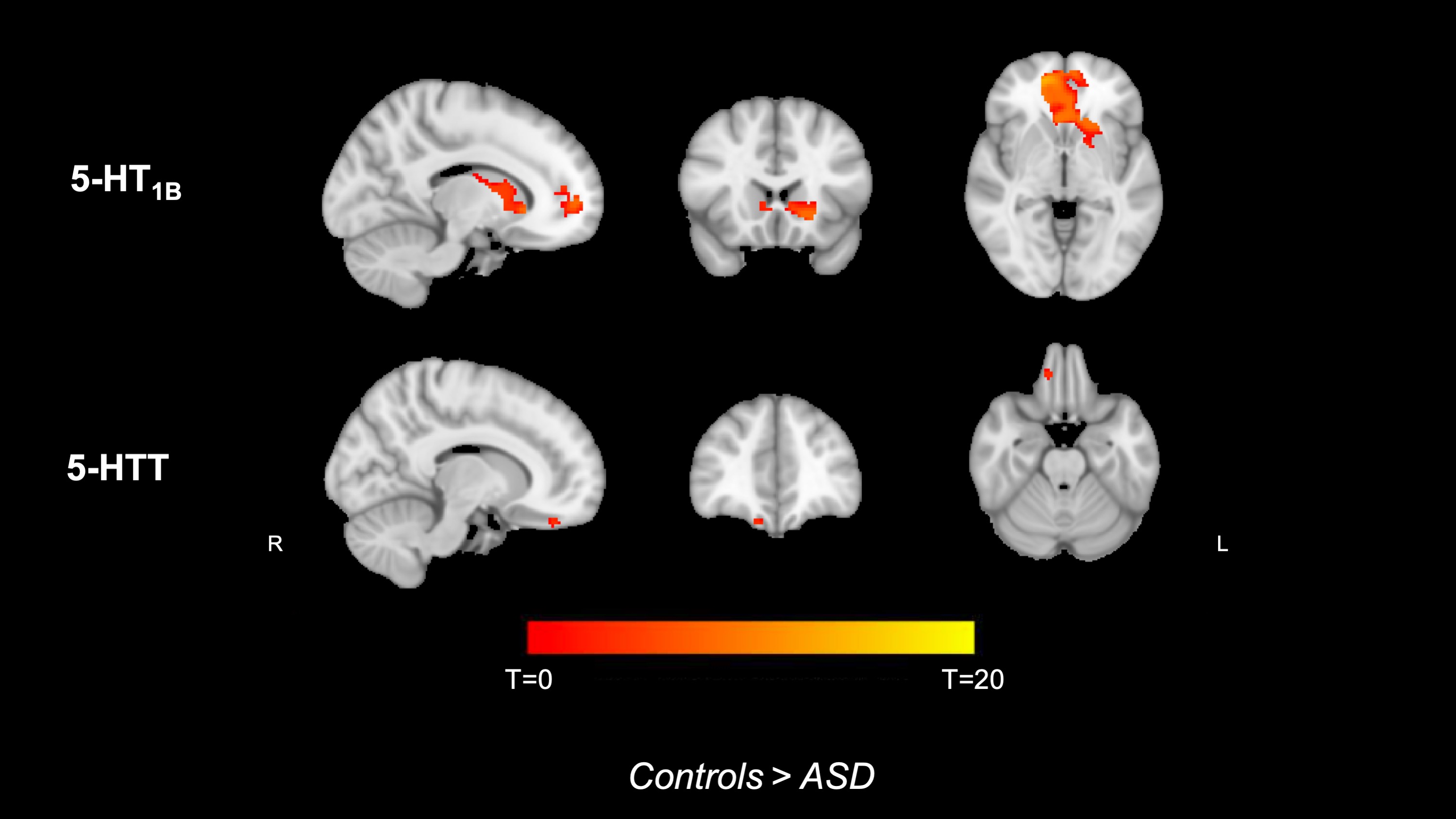

**Figure S2**. The serotonin 1B receptor (5-HT_1B_)-enriched and the serotonin transporter (5-HTT)-enriched Faces>Shapes functional response that are stronger in individuals without autism spectrum disorder (ASD) than that in individuals with ASD (highlighted red) at baseline placebo condition are overlaid on the anatomical brain. L=Left, R=Right.
